# Isolation and functional characterisation of lamina propria leukocytes from helminth-infected, murine small intestine

**DOI:** 10.1101/750786

**Authors:** Holly C. Webster, Anna T. Andrusaite, Amy L. Shergold, Simon W.F. Milling, Georgia Perona-Wright

## Abstract

The use of helminth infections as tools to understand the type 2 immune response is a well-established technique and important to many areas of immunological research. The phenotype and function of immune cell populations at the site of infection is a key determinant of pathogen clearance. However, infections with helminths such as the murine nematode *Heligomosmoides polygryrus* cause increased mucus production and thickening of the intestinal wall, which can result in extensive cell death when isolating and analysing cells from the lamina propria (LP). Populations of larger immune cells such as macrophages and dendritic cells are often trapped within mucus or dying tissues. Here we describe an optimised protocol for isolating LP leukocytes from the small intestine of *H.polygyrus* -infected mice, and we demonstrate phenotypic and functional identification of myeloid and CD4^+^ T cell subsets using cytokine staining and flow cytometry. Our protocol may provide a useful experimental method for the immunological analysis of the affected tissue site during helminth infections.

## Introduction

A type 2 immune response is established in response to stimuli such as allergens and helminth parasites. The type 2 immune response is characterised by infiltration of innate cells such as basophils, eosinophils and mast cells to mucosal sites, hyperplasia of goblet cells and a robust T helper 2 (Th2) cell response. The type 2 response is often associated with diseases such as allergy and asthma (Yazdanbakhsh et al., 2002), where the type 2 response is inappropriate. However, the type 2 immune response is critical for host protection during helminth infection and expulsion of parasites (Urban et al., 1991). Therefore, understanding the type 2 response and its regulation is important for fundamental science and for the development of new therapies. Murine helminth infections are widely used as models of type 2 immune activity, regulation and disease (Gause et al., 2003). A commonly used helminth is the nematode *Heligomosmoides polygyrus*, which naturally infects the small intestine of wild mice. This parasite is used as a model of human infection with hook worms such as *Necator americanus* and *Ancylostoma duodenale*, and as a model of a physiological Th2 response. *H. polygyrus* larvae are ingested orally at the L3 stage and, within 24 hours, reach the small intestine of their host and begin to migrate through the intestinal wall. Larvae migrate to the submucosa and encyst, undergoing two moults and developing into adult worms. Adult worms then migrate back through the intestinal wall and emerge into the lumen, where they mate and produce eggs (Monroy et al., 1992). The migration of larvae and adult worms through the small intestine wall results in epithelial cell damage. Damaged epithelial cells release cytokine alarmins such as IL-33, thymic stromal lymphopoietin (TSLP) and IL-25 (Barlow and McKenzie, 2011). IL-33 and IL-25 activate type 2 innate lymphoid cells (ILC2), which will in turn produce type 2 cytokines such as IL-5 and IL-13 (Barlow and McKenzie, 2011). Other innate cells including mast cells, eosinophils and basophils are rapidly recruited to the site of infection and also secrete IL-4, IL-5 and IL-13 (Davoine and Lacy, 2014; Gessner et al., 2005; Mukai et al., 2018). Macrophages and dendritic cells contribute to inflammation and stimulate the activation of Th2 cells, a subset of CD4^+^ effector T cells which are key to the type 2 immune response in helminth infection, allergy and asthma (Kim and Kim, 2018). Th2 cells produce the type 2 cytokines IL-5, IL-13 and IL-4. IL-4 drives further Th2 differentiation and guides class switching of B cells, while IL-5 and IL-13 are predominantly effector cytokines, recruiting and activating granulocytes, promoting degranulation, and guiding wound repair (Allen and Wynn, 2011; Liang et al., 2011) IL-13 stimulates goblet cell hyperplasia, which increases goblet cell production of mucin. Mucus production by goblet cells is a key component of helminth expulsion (Anthony et al., 2007).

One of the most common time-points investigated during *H.polygyrus* infection is fourteen days post infection, as Th2 cell expansion peaks (Perona-Wright et al., 2010), adult worms have emerged into the lumen (Hewitson et al., 2011) and both innate and adaptive immune responses are well established (Rolot and Dewals, 2018). At this time-point, there is large immune cell infiltration to the tissue, and further thickening occurs as a result of fibrosis. Mucus production is high due to goblet cell hyperplasia. Both factors contribute to extensive cell death, experimentally, when isolating leukocytes from the small intestine for further analysis. Published studies therefore typically concentrate on earlier time-points or on cells located in active lymphoid tissues (Mosconi et al., 2015; Pelly et al., 2016; Perona-Wright et al., 2010). Furthermore, larger immune cells such as macrophages frequently become trapped in mucus and dying tissues and are therefore very difficult to isolate. The ability to investigate immune cell populations at the site of infection is becoming increasingly important, as interactions between immune cells in inflamed tissues is emerging as key to fully understanding an immune response to infection. We have therefore optimised a method for isolating viable leukocytes from the small intestine LP during *H.polygyrus* infection and we show, using cytokine staining and flow cytometry, that this protocol enables the functional identification of subsets of both the myeloid and CD4^+^ T cell compartments.

## Methods

### Mice and Infection

Seven-week‐old female C57BL/6 mice were purchased from Envigo (Huntingdon, UK). Animals were maintained in individually ventilated cages under standard animal house conditions at the University of Glasgow and procedures were performed under a UK Home Office license (Project number 70/8483) in accordance with UK Home Office regulations following review by the University of Glasgow Ethics Committee. Mice were acclimatised for 1 week after arrival in the animal unit, and infected with 200 *H. polygyrus* L3 larvae by oral gavage.

### Isolation of lamina propria leukocytes

Naïve and infected animals were euthanised using carbon dioxide, and the small intestine removed by cutting below the stomach and above the caecum. Care was taken to ensure as much fat as possible was removed from the exterior of the intestine. Intestines were transferred immediately onto laboratory tissue paper soaked liberally in phosphate-buffered saline (PBS) and Peyer’s patches were removed using dissecting scissors, working quickly to ensure the tissue did not dry out. Continuing on PBS-soaked tissue paper and adding more PBS as needed to keep the tissue wet, the intestines were opened longitudinally using blunt tip scissors and then held by forceps and washed vigorously by agitation in a PBS-filled petri dish to remove faeces and mucus. Fine forceps were used to gently squeeze out any remaining mucus, running the forceps down the length of each intestine. Each intestine was then transferred onto a fresh piece of PBS soaked tissue and cut in to 1cm pieces. The pieces were collected and transferred to a 50ml centrifuge tube containing 30ml of HBSS (Gibco^™^ 14170088 no calcium, no magnesium) supplemented with 10% FCS (Gibco^™^ Fetal Bovine Serum, qualified, heat inactivated, E.U.-approved, South America Origin) and kept on ice. Each tube was shaken vigorously by hand and placed on ice for transfer from the animal unit back to the laboratory.

Further processing of the intestines was staggered, depending on sample number: a maximum of 8 samples was processed at one time and each step and wash was carried out quickly, ensuring that the intestinal tissue did not dry out at any stage in the protocol. Each sample was poured onto a large piece of 50-micron nitex mesh, folded into a funnel placed in a 400ml beaker. Samples were washed by pouring 30mls pre-warmed HBSS (Gibco^™^ 14170088 no calcium, no magnesium as before, but with no supplements), over the nitex mesh in the funnel. Using forceps, the samples were then transferred back into tubes containing 15ml 2mM EDTA (UltraPure^™^ 0.5M EDTA, pH 8.0 Cat. 15575020) in HBSS, pre-prepared and warmed to 37°C). The samples were shaken vigorously by hand and placed into an orbital shaker (Stuart, Orbital Incubator SI500) set to 220rpm and 37°C for 15 mins.

After shaking for 15min, the tubes were removed, the samples poured onto nitex mesh arranged in a funnel and washed with un-supplemented HBSS exactly as previously. After washing, the intestinal pieces were again transferred to tubes containing 15ml 2mM EDTA in HBSS and returned to the orbital shaker for a further 15 minutes. This process of EDTA washes was repeated twice more, for a total of three 15min shakes in 15ml 2mM EDTA in HBSS. After the 3^rd^ and final EDTA wash, the intestinal pieces were transferred into 15ml of RPMI 1640 (Gibco^™^ no glutamine, 21870076) supplemented with 10% FCS, 100 U/ml Penicillin Streptomycin (Life Technologies 15140122), 2mM L-glutamine (Life Technologies, 25030024) with 0.5mg/ml Collagenase VIII (Sigma-Aldrich C2139-500MG), all pre-prepared and warmed to 37°C. The samples were shaken vigorously by hand and placed back into the shaker set to 220rpm and 37°C. After 10 minutes, the samples were checked by removing from shaker and shaking vigorously by hand. The tissue had, as desired, began to break up and the media looked cloudy, reflecting cells released into the supernatant. The samples were placed back into shaker for a further 5 minutes, and then the visual check repeated. Optimal conditions meant that some but little tissue remained, and the supernatant was very cloudy. No sample was left in digestion buffer for any longer than 15 minutes, as this strongly reduced cell viability.

After the 15-minute digest period, enzyme action was stopped rapidly by adding 35ml of ice-cold RPMI 1640 containing 10% FCS, 1% Penicillin Streptomycin, 1% L-glutamine to each tube, and placing on ice. Each sample (all 50ml) was then filtered through a 100μm nylon mesh filter, followed by a 40μm nylon mesh filter, using a 25ml stripette and several 50ml centrifuge tubes. It was important not to crush through any tissue remaining on the top of the filters, as dying connective tissue decreased the viability of isolated cells. The samples were spun at 400g for 10 minutes at 4°C to pellet the isolated cells. The supernatants were discarded, and the pellets gently resuspended in 35ml of ice-cold RPMI 1640 (10% FCS, 1% Penicillin Streptomycin, 1% L-glutamine). The spin and resuspension of the pellet was repeated once more, as a final wash, and the resulting cell suspension was kept on ice, ready for further analysis.

### T cell cytokine stimulation and intracellular staining

Cells were resuspended at 3×10^6 cells/ml and 1ml of cell suspension was spun down at 400g for 5 minutes at 4°C in FACS tubes. Samples were resuspended in 500ul of RPMI 1640 supplemented with 10% FCS, 1% Penicillin Streptomycin and 1% L-glutamine and 2ul/ml of stimulation cocktail with protein inhibitors (Invitrogen eBioscience^™^ Cell Stimulation Cocktail plus protein transport inhibitors (500X)). Samples were incubated for 4 hours at 37°C in 5% CO2 incubator, and vortexed every hour. Samples were then washed twice with cold PBS before staining for flow cytometry. Dead cells were excluded using Fixable Viability Dye eFluor 780 (Ebioscience) and non-specific binding was blocked with Fc block anti-mouse CD16/32 Antibody (Clone 93, BioLegend). Samples were surface stained at 4°C for 20 min with: APC-conjugated anti-CD4 (RM4-5, BioLegend), FITC-conjugated anti-CD44 (IM7, BioLegend), and PerCpCy5.5-conjugated anti-TCRb (H57-597, BioLegend). Cells were fixed in 150ul of BD Cytofix/Cytoperm^™^ (554714) for 20 minutes at 4°C. Samples were then washed using BD Perm/Wash^™^ Buffer (554714) and 50ul of intracellular anti-cytokine antibody stain (PE-Cy7-conjugated anti-IL-13 (eBio13A, Invitrogen), PE-conjugated anti-IL-5 (TRFK5, BioLegend), e450-conjugated anti-IFNy (XMG1.2, Invitrogen)) or appropriate isotype control added to each sample and incubated at room temperature, protected from light, for 1 hour. Samples were washed using BD Perm/Wash^™^ Buffer and acquired immediately on the BD LSRII flow cytometer running FACS-Diva software (BD Biosciences). Analysis was performed using FlowJo (Treestar).

### Myeloid cell staining and gating

For myeloid cell staining, dead cells were excluded using Fixable Viability Dye eFluor 780 (Ebioscience) and non-specific binding was blocked with Fc block anti-mouse CD16/32 Antibody (Clone 93, BioLegend). Samples were then surface stained at 4°C for 30 min with the following mAbs: AlexaFluor 647-conjugated anti-CD3 (17A2, BioLegend), AlexaFluor 647-conjugated anti-B220 (RA3-6B2, BioLegend), AlexaFluor 700-conjugated anti-MHCII (M5/114.15.2,BioLegend), PerCP-Cy5.5-conjugated anti-CD11c (N418, BioLegend), BV605-conjugated anti-CD11b (M1/70, BioLegend), BV510-conjugated anti-Ly6C (HK1.4, BioLegend), PE-conjugated anti-Ly6G (1A8, BioLegend), and PE-Cy7-conjugated anti-CD64 (X54-5/7.1, BioLegend). Cells were then washed with ice cold PBS (supplemented with 10% FCS and 2mM EDTA) at 400g for 10 minutes at 4°C and acquired immediately on BD Fortessa flow cytometer running FACS-Diva software (BD Bioscience). Analysis was performed using FlowJo (Treestar).

## Results

Several protocols exist for the isolation of leukocytes from the small intestine of mice, but the inflamed and mucus-rich conditions of helminth-infected intestines have proven challenging. We set out to optimise isolation conditions for leukocytes from the small intestine LP of mice 14 days post-infection with *H.polygyrus*, adapting protocols previously used by the Milling group for naïve and bacterially-infected tissues (Bravo-Blas et al., 2019; Cerovic et al., 2013). Our previous protocols had resulted in extensive cell death when processing *H. polygyrus* infected tissues, which we reduced by optimisation of the dissection and digestion workflow and the concentrations of digestion enzymes and EDTA solutions (**Figure 1**). We also tried the inclusion of dithiothreitol (DTT) to disrupt mucus, and the use of a density gradient to remove dead cells (Goodyear et al.), but we found that DTT reduced cell viability and the density gradient reduced cell yield. Our optimised protocol removes both steps. When assessing the forward and side scatter of the isolated leukocytes, our optimised protocol showed clear lymphocyte populations that were not present in our previous isolations (**Figure 1A**). Furthermore, when comparing the viability between these experiments, cells isolated using our optimised protocol had much higher viability compared to previous experiments (**Figure 1B and 1C**).

**Figure 1.**
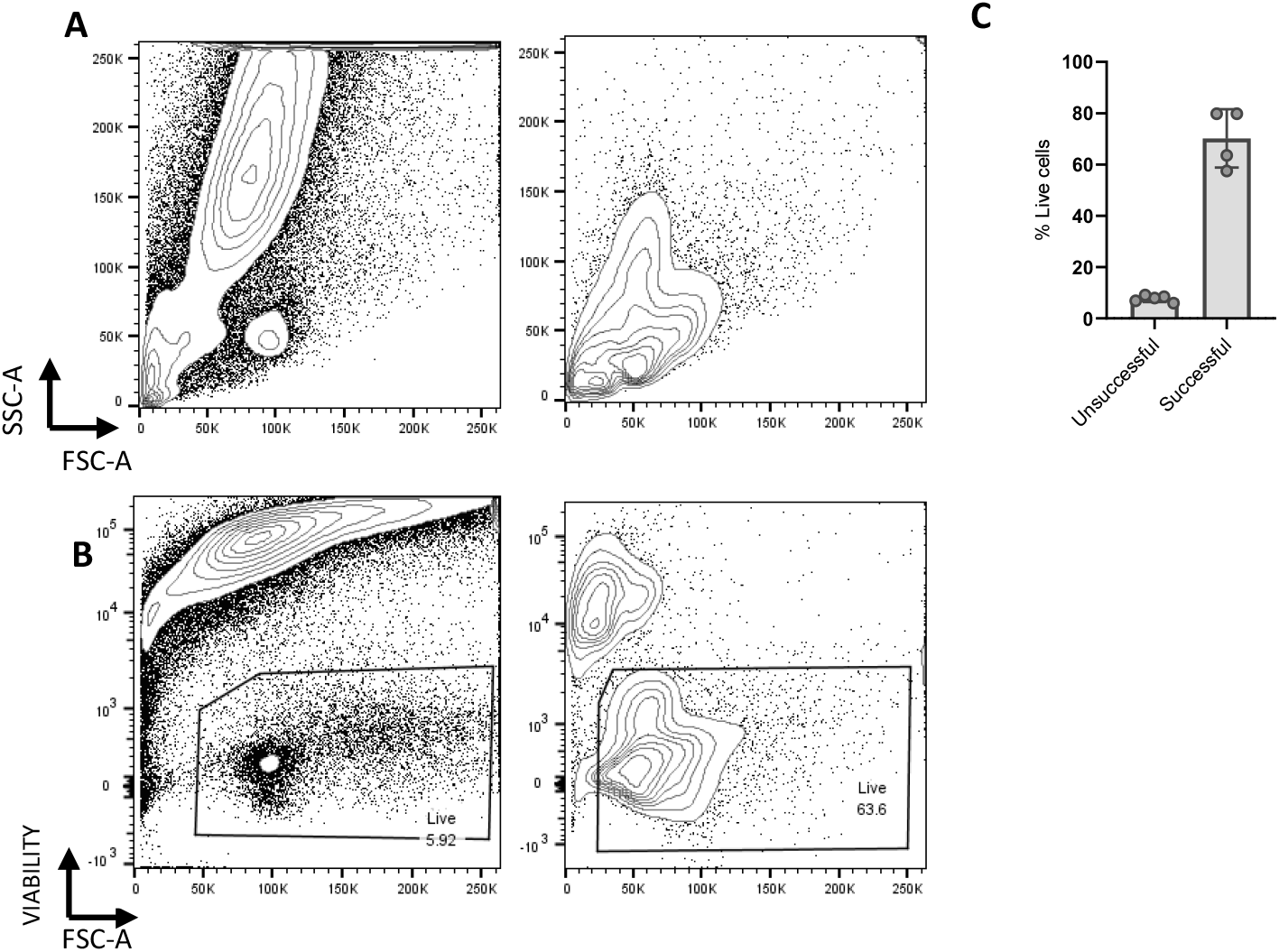
Scatter and viability of unsuccessful vs successful isolation protocols. C57BL/6 mice were infected with 200 L3 *H.polygyrus*. 14 days later the small intestine was removed, and leukocytes isolated and analysed by flow cytometry. (**A**) Representative scatter of unsuccessful (left) and successful (right) protocols. (**B**) Representative viability of unsuccessful (left) and successful (right) protocols. Data are representative of 3 separate experiments with n=4−5 in each experiment.

We next assessed the effectiveness of our protocol in the isolation of CD4^+^ T cells at different times after *H.polygyrus* infection (**Figure 2**). The small intestines of naïve mice or mice 7 and 14 days post-infection with *H.polygyrus* were collected, leukocytes were isolated and CD4^+^ T cells subsequently analysed by flow cytometry. Live T cells were recovered at all times after infection (**Figure 2A-C**), and the total number of recovered CD4+ T cells showed a trend towards higher yields from infected animals than in naïve, and from day 14 than day 7 (**Figure 2D**). The viability of cells remained largely consistent between different timepoints, with a slight decrease at day 14 post-infection (**Figure 2E**).

**Figure 2.**
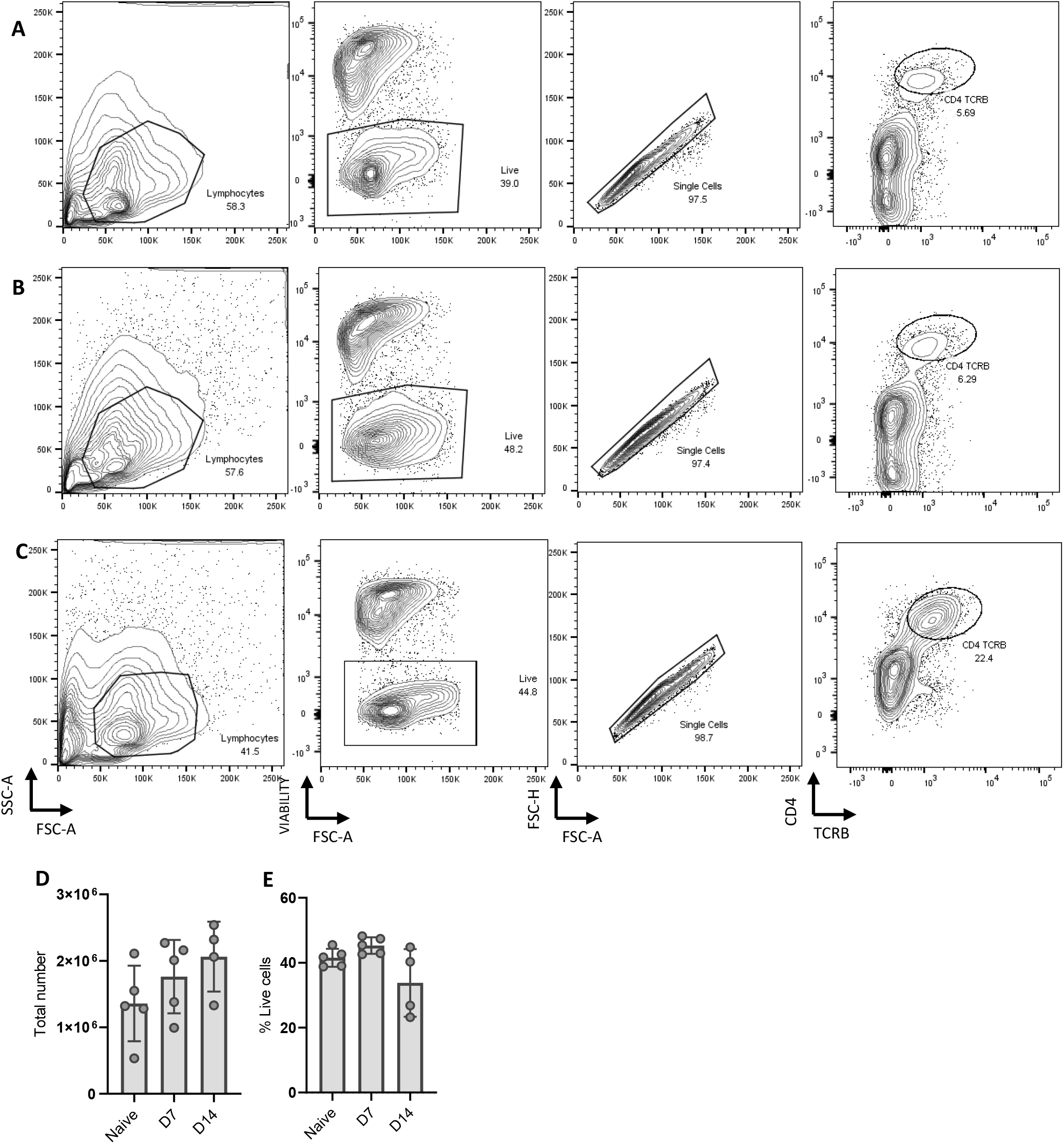
Timepoint isolation of CD4+ T cells from the small intestine of *H.polygyrus* infected mice. C57BL/6 mice were uninfected or infected with 200 L3 *H.polygyrus* larvae. 7 and 14 days later, the small intestine was removed, and leukocytes isolated and analysed by flow cytometry. Representative CD4+ T cell gating strategy of (**A**) naïve (**B**) 7 days post-infection and (**C**) 14 days post-infection. (**D**) The total number of CD4+ T cells at each timepoint (**E**) The percentage of live cells in the lymphocyte gate at each timepoint. Data are representative of 3 separate experiments with n=4−5 in each experiment.

To assess the function of recovered CD4^+^ T cells, we next examined effector subsets based on cytokine secretion. This technique requires a 4-hour stimulation with PMA and ionomycin in the presence of Golgi inhibitors. As this is a harsh stimulation, it often decreases cell viability even in cells from easily accessible tissues and, in our previous isolation protocols, cells isolated from the small intestine of *H.polygyrus* infected mice showed almost complete cell death after stimulation. To test the current method, we stimulated bulk cells and stained intracellularly for the type 2 cytokines IL-5 and IL-13 (**Figure 3**). Live cells were recovered, and cytokines were detected at all time points post-infection. As predicted, IL-5 and IL-13 positive cells were more frequent in intestinal samples from infected mice compared to those from naïve controls (**Figure 3A-C, D and E**).

**Figure 3.**
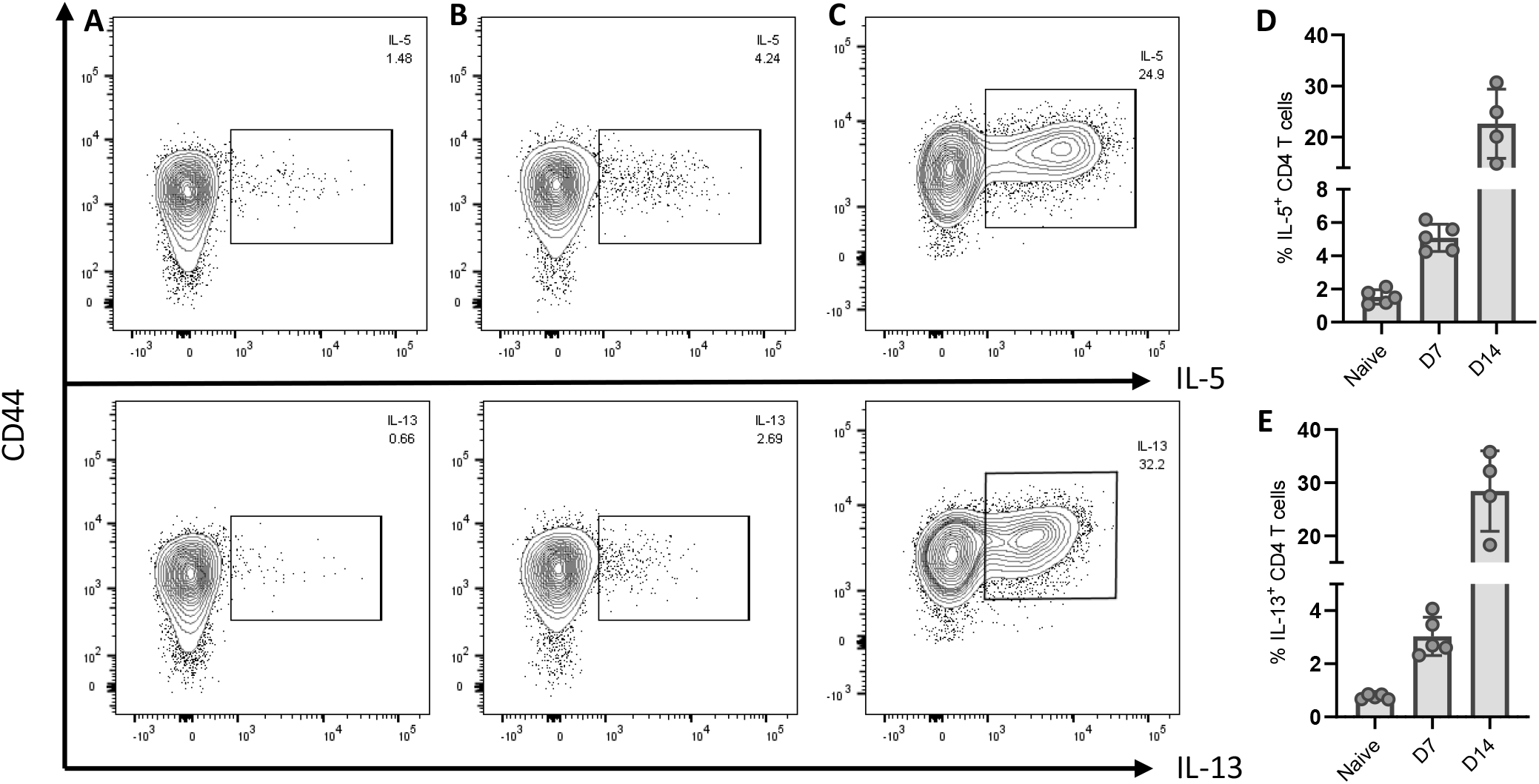
Type 2 cytokine secretion by CD4^+^ T cells during *H.polygyrus* infection. C57BL/6 mice were uninfected or infected with 200 L3 *H.polygyrus* larvae. 7 and 14 days later the small intestine removed, and leukocytes isolated, and intracellular cytokines analysed by flow cytometry. Representative CD4+ T cytokine secretion of IL-13 (right) and IL-5 (left) of (**A**) naïve mice (**B**) 7 days post-infection and (**C**) 14 days post-infection. (**D**) Percentage of IL-5^+^ and (**E**) IL-13^+^ CD4^+^ T cells at each timepoint. Data representative if 3 separate experiments and n=4/5.

Isolating larger immune cells such as myeloid cells from *H.polygyrus* infected small intestines has also been difficult previously. These cells are typically less robust than CD4^+^ T cells in surviving the isolation process, and the extent of tissue digestion must be finely balanced between sufficient digestion to release adherent cells but gentle enough that cell viability is preserved. We therefore optimised our protocol also to examine the myeloid compartment in the small intestine at different times after *H. polygyrus* infection (**Figure 4**).

**Figure 4.**
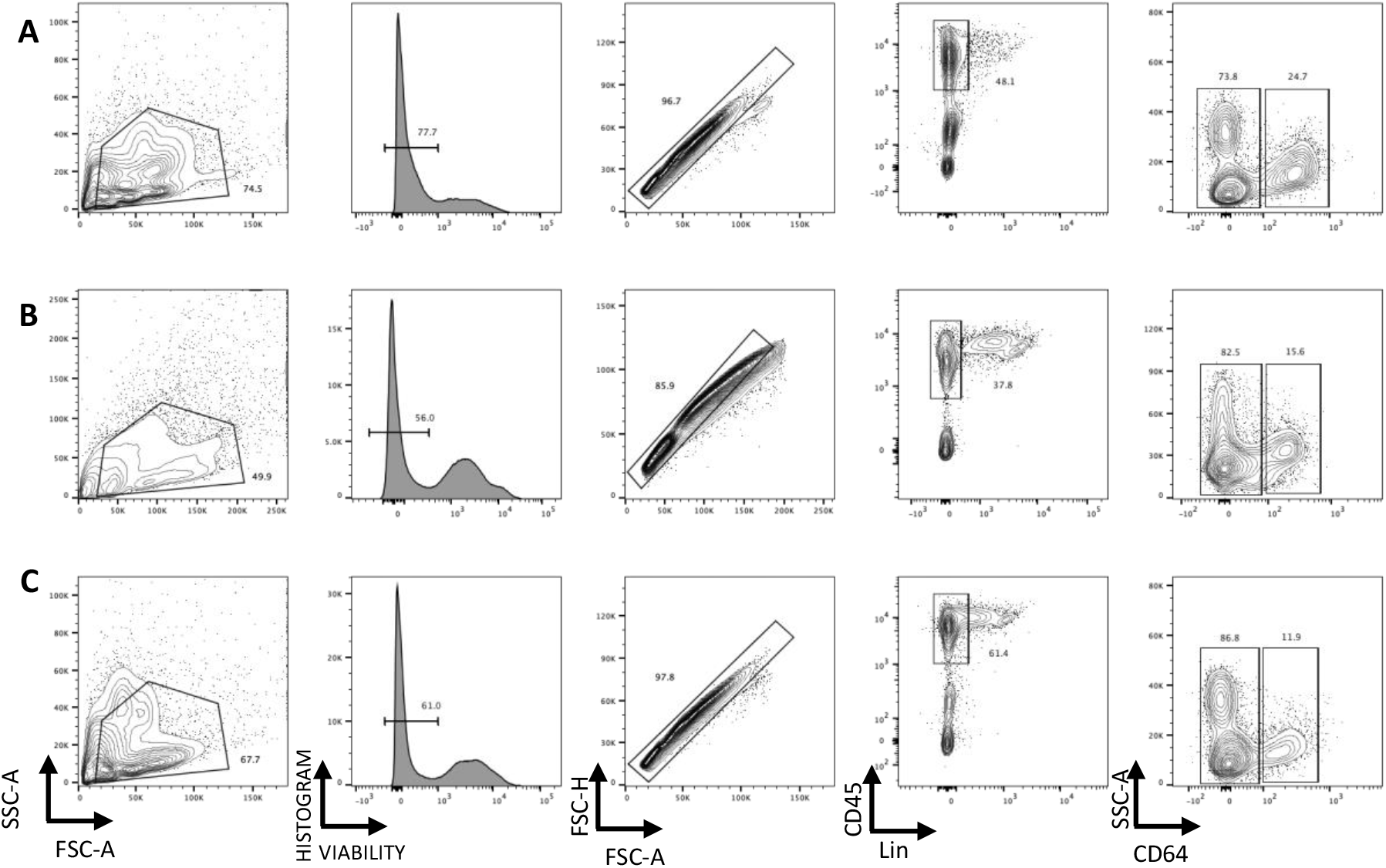
Isolation of myeloid cells from the small intestine of *H.polygyrus* infected mice. C57BL/6 mice were uninfected or infected with 200 L3 *H.polygyrus* larvae and 7 and 14 days later the small intestine removed and leukocytes isolated and analysed by flow cytometry. Representative myeloid cell gating strategy of naïve (**A**) 7 days post-infection (**B**) and 14 days post-infection (**C**). Data representative of 3 separate experiments.

The isolated leukocytes contained a substantial live cell population that is CD45^+^ but negative for T cell and B cell lineage markers. Using CD64 expression as a marker of monocyte/macrophages that distinguishes these cells from dendritic cell populations in the murine intestine (Cerovic et al., 2014), we confirmed that our protocol can isolate live myeloid cell populations at 14 days post *H. polygyrus* infection (**Figure 4**). We also recovered a sufficient number of cells from the infected LP to further analyse distinct subsets of the myeloid lineage (**Figure 5**). Following the gating illustrated in Figure 4 and selecting CD64^+^ cells, we identified infiltrating monocytes (MHCII^−^, Ly6C^+^), maturing monocytes (MHCII^+^, Ly6C^+^) and macrophages (MHCII^+^, Ly6C^−^) (**Figure 5A**), illustrated in the ‘monocyte waterfall’ (Bain et al., 2014), at both day 7 and day 14 post infection. Similarly, we identified infiltrating neutrophils (live single CD45^+^ Lin^−^ CD64^−^ Ly6G^+^ CD11b^+^ cells) (**Figure 5B**). Finally, we are also able to distinguish dendritic cell populations by gating on live single CD45^+^ Lin^−^ CD64^−^ CD11c^+^ MHCII^+^ cells (**Figure 5C**). Using these gating strategies,we calculated the absolute number of these myeloid cells within the small intestine LP (**Figure 6**). Monocytes (MHCII^−^, Ly6C^+^) and maturing monocytes (MHCII^+^, Ly6C^+^) showed a clear increase in number over the course of *H. polygyrus* infection (**Figure 6A**). Similarly, the abundance of neutrophils (**Figure 6B**) and dendritic cells (**Figure 6C**) also significantly increased at day 7 of infection but appeared to subside by day 14. Together these results confirm that the method described here yields high quality cell samples from *H. polygyrus* infected intestinal tissue, that can be further analysed for detailed immune cell subsetting and characterisation.

**Figure 5.**
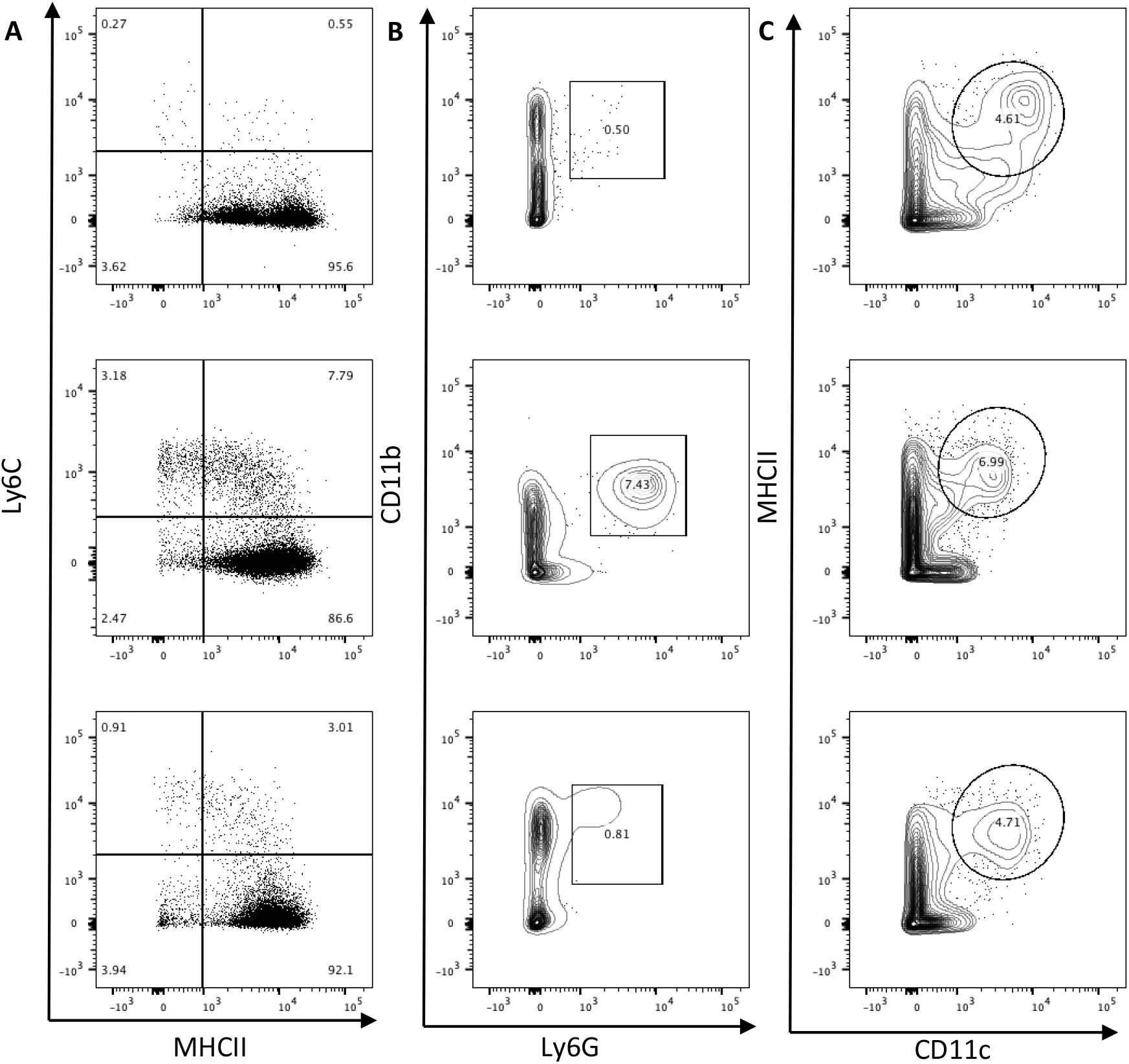
Isolation of monocytes, macrophages, neutrophils and dendritic cells from small intestinal LP during H.polygyrus infection. C57BL/6 mice were uninfected (left) or infected with 200 L3 *H.polygyrus* larvae and 7 (middle) and 14 (left) days later the small intestine removed and leukocytes isolated and analysed by flow cytometry. (**A**) Monocytes (MHCII^−^, Ly6C^+^), maturing monocytes (MHCII^+^, Ly6C^+^) and macrophages (MHCII^+^, Ly6C^−^) isolated at different time points of infection. Gated on live, single, CD45^+^, Lin^−^, CD64^+^ cells. (**B**) Neutrophils (Ly6G^+^, CD11b^+^) isolated at different time points of infection. Gated on live, single, CD45+, Lin-, CD64-cells. (**C**) Dendritic cells (MHCII^+^, CD11c^+^) isolated at different time points of infection. Gated on live, single, CD45^+^, Lin^−^, CD64^−^ cells. Data representative of 3 separate experiments.

**Figure 6.**
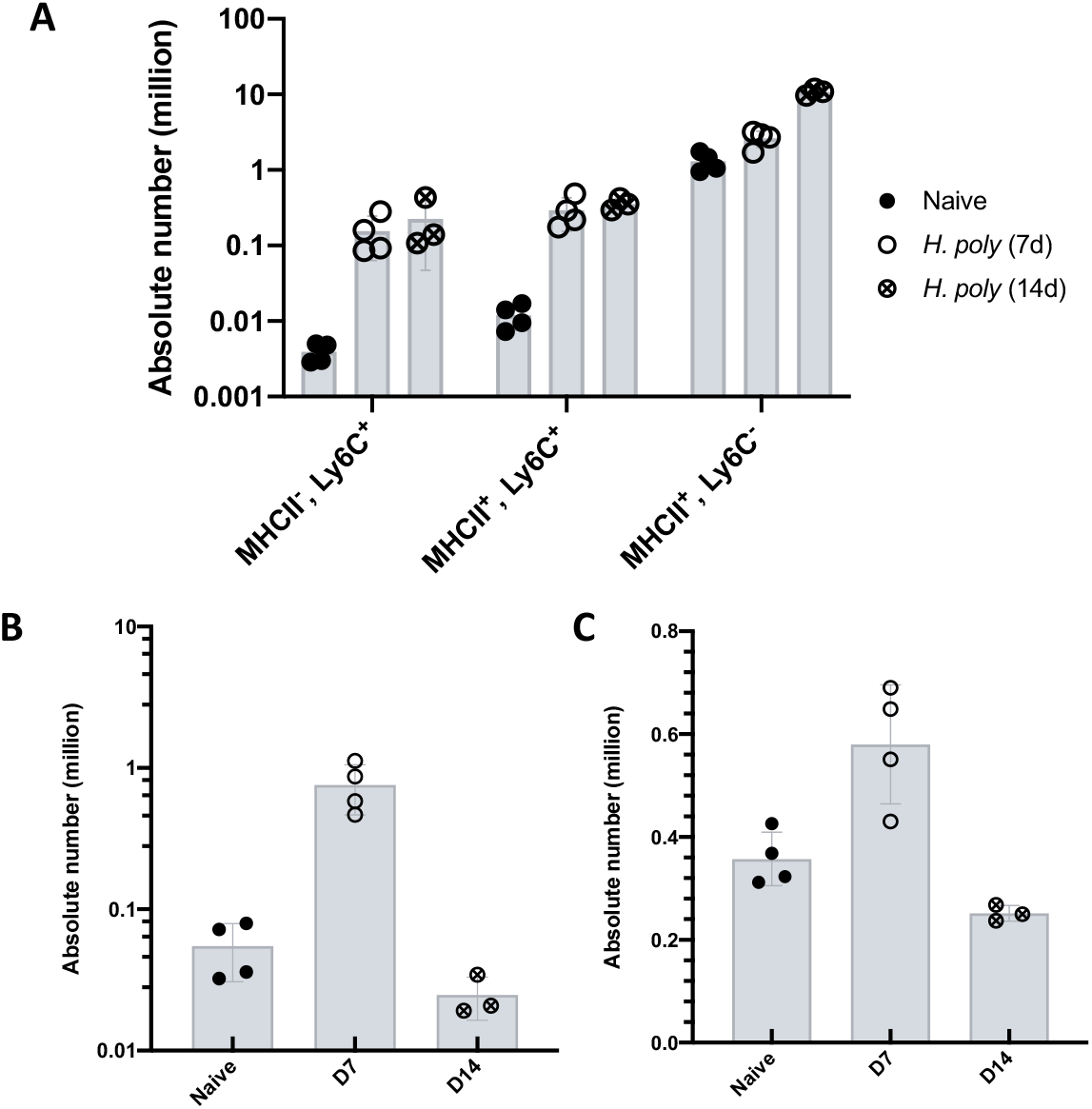
Absolute number of monocytes, macrophages, neutrophils and dendritic cells recovered from small intestinal LP during *H.polygyrus* infection. C57BL/6 mice were uninfected or infected with 200 L3 H.polygyrus larvae and 7 and 14 days later the small intestine removed, and leukocytes isolated and analysed by flow cytometry. (A) Absolute number of monocytes (MHCII^−^,Ly6C^+^), maturing monocytes (MHCII^+^, Ly6C^+^) and macrophages (MHCII^+^, Ly6C^−^) isolated at different time points of infection. Gated on live, single, CD45^+^, Lin^−^, CD64^+^ cells. (B) Absolute number of neutrophils (Ly6G^+^, CD11b^+^) isolated at different time points of infection. Gated on live, single, CD45^+^, Lin^−^, CD64^−^ cells. (C) Absolute number of dendritic cells (MHCII^+^, CD11c^+^) isolated at different time points of infection. Gated on live, single, CD45^+^, Lin^−^, CD64^−^ cells. Data representative of 2 separate experiments (n=3/4).

## Discussion

Our data demonstrate that we have optimised a protocol that allows for successful isolation and functional characterisation of leukocytes from the small intestine of *H.polygyrus* infected mice at two key timepoints, day 7 and day 14 post-infection. We were able to identify a clear CD4^+^ Th2 cell population whose frequency and number increased in infection, reflecting the predicted immunobiology of the infection and confirming that, even at day 14 when mucus levels are highly elevated, this protocol is able to recover representative cell populations. Furthermore, we were able to assess cytokine secretion by CD4^+^ T cells from the infected tissue and maintain cell viability of more than 35% despite re-stimulation of these cells, which previously had resulted in extensive cell death. We also demonstrate that we can isolate distinct populations of monocytes, macrophages, neutrophils and dendritic cells from the small intestine LP of helminth-infected animals, which had previously been a particular challenge due to the fragile nature of myeloid cells *ex-vivo*. Together, our data indicate that the isolation protocol that we present here allows the phenotypic and functional characterisation of innate and adaptive immune cells active in the infected tissue site during a Type 2 immune response to helminth infection.

The isolation of viable leukocytes from the intestinal LP of naïve mice or those infected with bacterial, viral and protozoan pathogens has been widely published (Bravo-Blas et al., 2019; Cerovic et al., 2013; Goodyear et al., 2014; Isakov et al., 2011; Perona-Wright et al., 2012). In contrast, helminth-infected tissues have been difficult to use as sources of immune cells. This is largely attributed to the ‘weep and sweep’ effector mechanisms associated with a type 2 immune response, including extensive mucus production, activation of pro-fibrotic macrophages, lymphocytic and granulocytic infiltration, and the release of histamine and cytokines that lead to a mobile and fragile intestinal epithelium (Allen and Maizels, 2011; Allen and Wynn, 2011; Anthony et al., 2007; Cliffe et al., 2005; Webb and Tait Wojno, 2017). Our understanding of type 2 immunity has therefore been biased towards observations made in the early stages of infection or in lymphoid tissues. Here, we have optimised the timing, the digestion and the processing of helminth-infected intestinal tissue and we present a technique that allows immune characterisation of the small intestine lamina propria during an acute type 2 immune response. By aiming to remove rather than add steps to the isolation protocol, we chose to prioritise speed over precision and were able to achieve sample viabilities ranging from 35-75% of lymphocytes, even after stimulation with PMA and ionomycin (**Figures 2 and 3**). A danger of selecting a light approach was that we would lose cells that were more firmly attached in the inflamed tissue. However, our analysis of myeloid populations (**Figures 4**, **5 and 6**) indicates that these cells are also isolated and viable following this protocol. We therefore present a new method suitable for the analysis of both myeloid and lymphoid cells in the lamina propria of mice 14 days after *H.polygyrus* infection.

There is growing interest in the immunobiology of infections at the effector site, rather than associated lymphoid tissues. The characterisation of CD4^+^ and CD8^+^ resident memory T cells in the affected tissues has redefined our models of T cell memory (Carbone and Gebhardt, 2019). T cell cytokines are thought to be segregated by tissue, with IL-4 concentrated in the active lymph node and IL-5 and IL-13 dominating in the infected tissue (Liang et al., 2011; Redpath et al., 2015). Myeloid cells in the effector site are key to tissue remodelling, clearing of microorganisms and debris, and the initiation and perpetuation of appropriate T cell responses (Allen and Wynn, 2011; Kim and Kim, 2018; Rolot and Dewals, 2018). Being able to examine cells and their activation at the tissue site will be critical for understanding the dynamics of the *H. polygyrus* infection model as well as those of a prototypical type 2 immune response. We therefore present a protocol to allow for successful analysis of leukocyte populations at the site of *H.polygyrus* infection, the small intestine LP, throughout the course of infection.

## Acknowledgements

We thank David Dow for animal husbandry; Nicola Britton, Claire Ciancia and Rick Maizels for *H.polygyrus* larvae and life cycle maintenance, and for advice and discussion; the Flow Core at the University of Glasgow and especially Diane Vaughan for flow cytometry support; and Alberto Bravo-Blas and Graham Heieis for early development of isolation and staining protocols. This work was supported by the MRC (grant MR/S009779/1) and funds from the University of Glasgow. HCW and AA are PhD scholars supported by the University of Glasgow MVLS PhD studentship programme.

